# Ultramutagenesis through combined Msh2 depletion and proofreading-deficient DNA polymerase ε expression enhances small molecule target identification in forward genetic screens

**DOI:** 10.64898/2026.06.26.734921

**Authors:** Alexander-Hoi Nguyen, Vicky Li, Wei-Min Chen, Jacob Kimberg, Anthony J. Davis, David G. McFadden, Juan Manuel Povedano

## Abstract

DNA mismatch repair (MMR) deficiency has been widely utilized in forward genetic screens to identify drug resistance mutations and elucidate the mechanisms of action of cytotoxic small molecules. However, MMR deficiency generates a characteristic mutational signature enriched for C>T transitions that precludes saturation of mutagenesis. We hypothesized that expression of proofreading deficient DNA polymerase epsilon, as has been observed in patients with biallelic MMR deficiency, would enhance mutagenesis and enable identification of new resistance mutations and cytotoxin mechanisms. Here, we combine auxin-inducible degradation of Msh2 with doxycycline-inducible expression of the proofreading-deficient Polε^P286R^ mutant in murine small cell lung cancer (SCLC) cells. Simultaneous MMR deficiency and mutant Polε expression markedly increased the emergence of bortezomib-resistant clones relative to either perturbation alone and targeted sequencing of *Psmb5* in bortezomib-resistant clones identified recurrent mutations previously reported in chemical mutagenesis-based resistance screens. We next applied this platform to investigate resistance to lurbinectedin, a clinically relevant therapy for SCLC. Whole-exome sequencing revealed recurrent loss-of-function alterations in nucleotide excision repair (NER) genes, most notably *Ercc5* and *Ercc4*, implicating NER deficiency as a major mechanism of lurbinectedin resistance and supporting a central role for NER in mediating lurbinectedin-induced cytotoxicity. Collectively, these findings establish inducible ultramutagenesis as a powerful and versatile platform for the unbiased discovery of drug resistance mechanisms and therapeutic vulnerabilities.

## INTRODUCTION

The identification of the molecular mechanism of action of cytotoxic small molecules is a critical step in drug development. Knowledge of the direct target of a small molecule provides insight into target engagement, on-target effects, and potential mechanisms of resistance. Genetic approaches to elucidate the mechanism of action commonly rely on the identification of mutations that confer resistance by disrupting the interaction between a small molecule and its protein target^1, 2^. Such strategies require the generation of sufficient genetic diversity to enable the emergence and selection for resistant clones. Historically, elevated mutation rates have been achieved through chemical mutagenesis or by utilizing cell lines with defects in mismatch repair (MMR), which exhibit increased spontaneous mutagenesis ^2–5^. These approaches have been successfully employed to identify resistance-conferring alleles and uncover the molecular targets of cytotoxic compounds. However, the spectrum of mutations generated by MMR deficiency is biased, potentially limiting the identification of resistance.

One approach to generating MMR-deficient cancer cell lines is through genetic inactivation of *MSH2* or the use of inducible protein degradation systems targeting essential MMR components^5, 6^. We and others have previously demonstrated that MMR deficiency facilitates the acquisition of resistance-conferring mutations to a variety of cytotoxic small molecules ^5–10^. Despite these successes, the published literature likely overestimates the overall efficiency of this strategy. In our experience, only a subset of small molecules yields resistant clones in MMR-deficient cells, highlighting important limitations of MMR-based mutagenesis approaches. Several factors may contribute to the low success rate. First, some small molecules may not exert their effects through a discrete protein target, thereby hindering the emergence of target-based resistant mutations. Second, resistance-conferring mutations must disrupt compound activity while preserving the essential physiological function of the target protein, a constraint that may substantially limit the number of viable resistance alleles in essential proteins. Third, the mutational spectrum generated by MMR deficiency is enriched for C>T transitions and may not generate the full spectrum of potential resistance mutations.

Biallelic mismatch repair deficiency (bMMRD) is a rare hereditary cancer predisposition syndrome caused by germline loss-of-function mutations in core MMR genes. Defective MMR promotes the rapid accumulation of somatic mutations and frequently leads to the acquisition of secondary mutations in the proofreading (exonuclease) domains of DNA polymerases ε (Polε) and δ (Polδ). The combined impairment of DNA repair and replication fidelity drives exceptionally high mutation rates, termed “ultramutation,” resulting in tumors with mutation frequencies that exceed 250 mutations per megabase^11^. This natural occurring synergy between MMR deficiency and defective polymerase proofreading provides a compelling model for engineering extreme mutator phenotypes.

In this study, we engineered inducible systems for Msh2 degradation and expression of a proofreading-deficient Polε mutant into cancer cell lines to enhance the spectrum of mutations capable of generating small molecule resistance. Using this platform, we screened for resistance to two cytotoxic small molecules and successfully identified recurrent mutations in genes encoding proteins required for their cytotoxic activity. These findings establish inducible ultramutagenesis as an effective strategy for forward genetic screens aimed at elucidating drug mechanisms of action and resistance pathways.

## RESULTS

We hypothesized that combination of MMRd and expression of exonuclease deficient Polε would increase and broaden the spectrum of mutations in cancer cell lines. To generate an inducible MMR-deficient model, we used CRISPR-Cas9 to engineer an auxin-inducible degron (AID) tag into the endogenous *Msh2* locus of murine SCLC (518T2) cells (Figure 1A)^12^. Following blasticidin selection, single-cell clones were isolated, expanded, and screened for *Msh2* expression by western blot analysis (Figure 1B). We identified a clone that lacked detectable wild-type Msh2 protein and instead expressed a higher-molecular-weight species consistent with successful integration of the AID tag at the endogenous *Msh2* locus (Figure 1B).

**Figure 1.**
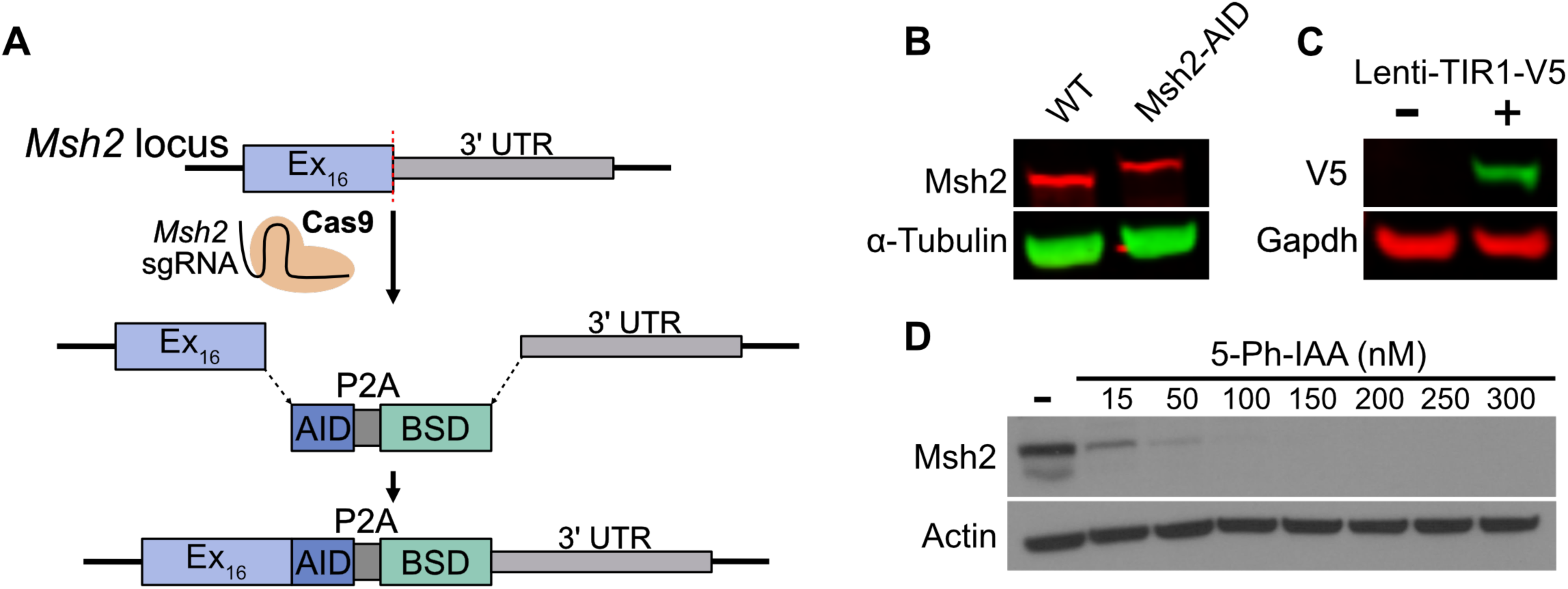
Engineering an auxine-inducible degron into the Msh2 locus in murine SCLC cells. (A) Schematic depicting the knock-in of an AID tag followed by a P2A and blasticidin resistant cassette (BSD). (B) Western-blot showing the expression of Msh2 (left) and the MSh2-AID (right) from murine SCLC cells and engineeered Msh2-AID cells, respectively. α-Tubulin was used as loading control. (C) Western-blot of lentiviral transduced murine SCLC cells showing the expression of the plant Cullin adapter protein, TIR1, using an antibody recognizing the V5 tag. Gapdh was used as loading control. (D) Western-blot showing the degradation of Msh2-AID in a dose-dependent manner at 24h. Actin was used as loading control.

To enable inducible degradation of AID-tagged Msh2, we introduced the auxin-responsive E3 ubiquitin ligase adaptor protein TIR1^F74A^ via lentiviral transduction and confirmed stable expression by immunoblot analysis using a V5 antibody (Figure 1C and Supplementary Figure 1). The TIR1^F74A^ variant exhibits enhanced affinity for the synthetic auxin analog 5-phenyl-indole-3-acetic acid (5-Ph-IAA), allowing efficient target protein degradation at substantially lower concentrations^13, 14^. Treatment of Msh2-AID;TIR1^F74A^ cells with 150-200 nM 5-Ph-IAA induced rapid and efficient degradation of Msh2, with complete loss of detectable protein observed within 24 hours (Figure 1D). Collectively, these results established a robust and tunable system for the inducible degradation of endogenous Msh2 in murine SCLC cells, enabling temporal control of MMR deficiency.

Next, we used a PiggyBac transposon-based vector to introduce a *Polε* transgene harboring the P286R exonuclease-domain mutation, which impairs proofreading activity and increases replication-associated mutagenesis^15^. The transgene was placed under the control of a doxycycline-inducible Tet-On system containing a TRE3G promoter, enabling temporal regulation of *Polε*^P286R^ expression (Figure 2A). Treatment of 518T2 cells with non-toxic concentrations of doxycycline resulted in robust induction of *Polε*^P286R^ expression within 48 hours, as confirmed by immunoblot analysis (Figure 2B and Supplementary Figure 2). These engineered cells enabled independent or combined manipulation of two complementary mutagenic pathways: (1) induction of MMR deficiency through Msh2 degradation following 5-Ph-IAA treatment, (2) expression of proofreading-deficient *Polε*^P286R^ following doxycycline treatment, (3) simultaneous induction of both perturbations through combined treatment with 5-Ph-IAA and doxycycline (Figure 3A). This experimental system provided a versatile platform to evaluate the individual and combined contributions of MMR deficiency and defective polymerase proofreading to mutagenesis and the emergence of drug-resistant clones.

**Figure 2.**
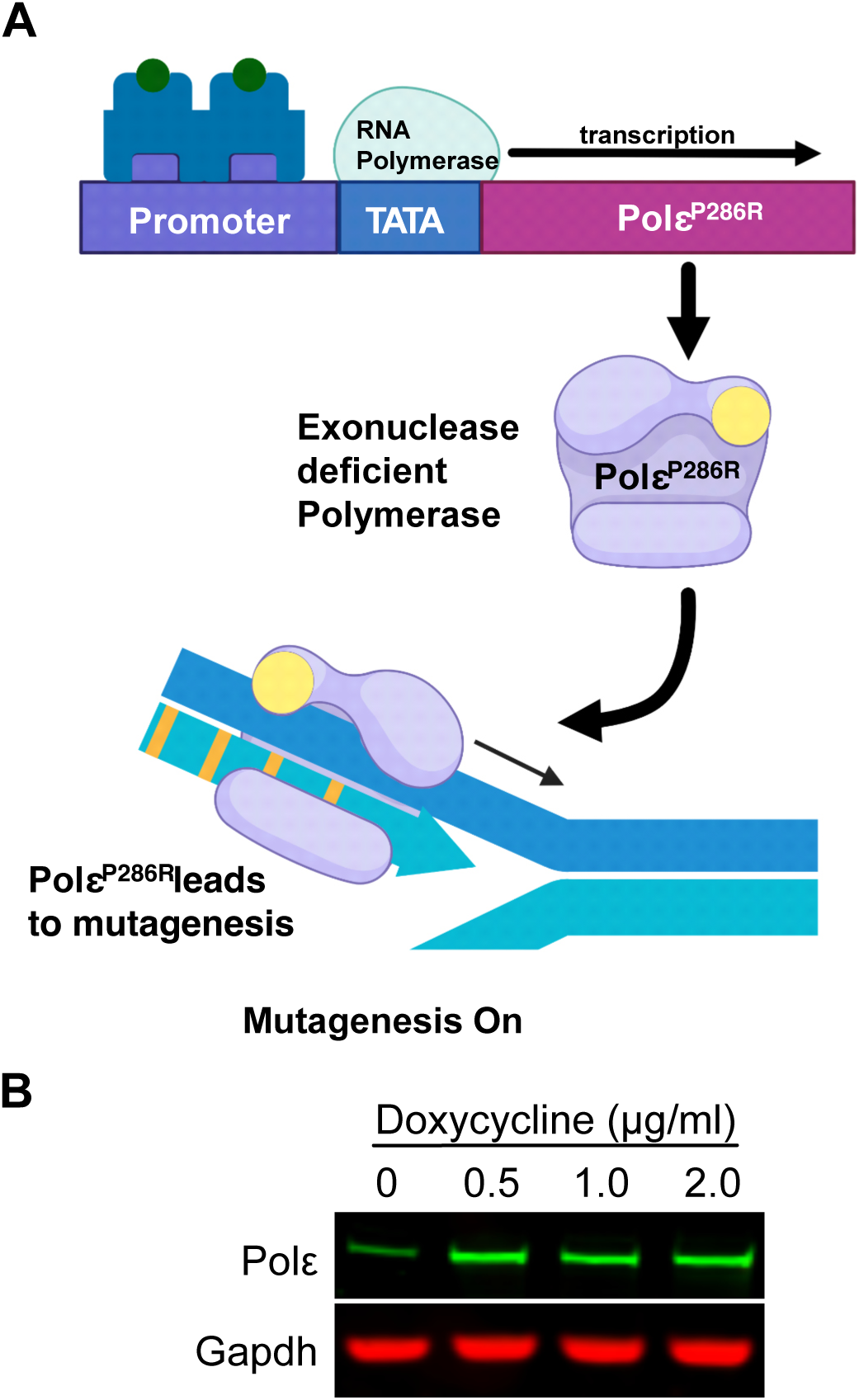
Tetracycline inducible promoter for mutant Polɛ overexpression in murine SCLC cells. (A) Schematic depicting the tetracycline-inducible expression of catalytically dead polymerase ε, leading to mutagenesis. (B) Western-blot showing the expression of Polε in a dose-dependent manner at 48 hours in TREG-PolεP286R SCLC cells. Gapdh used as a loading control.

**Figure 3.**
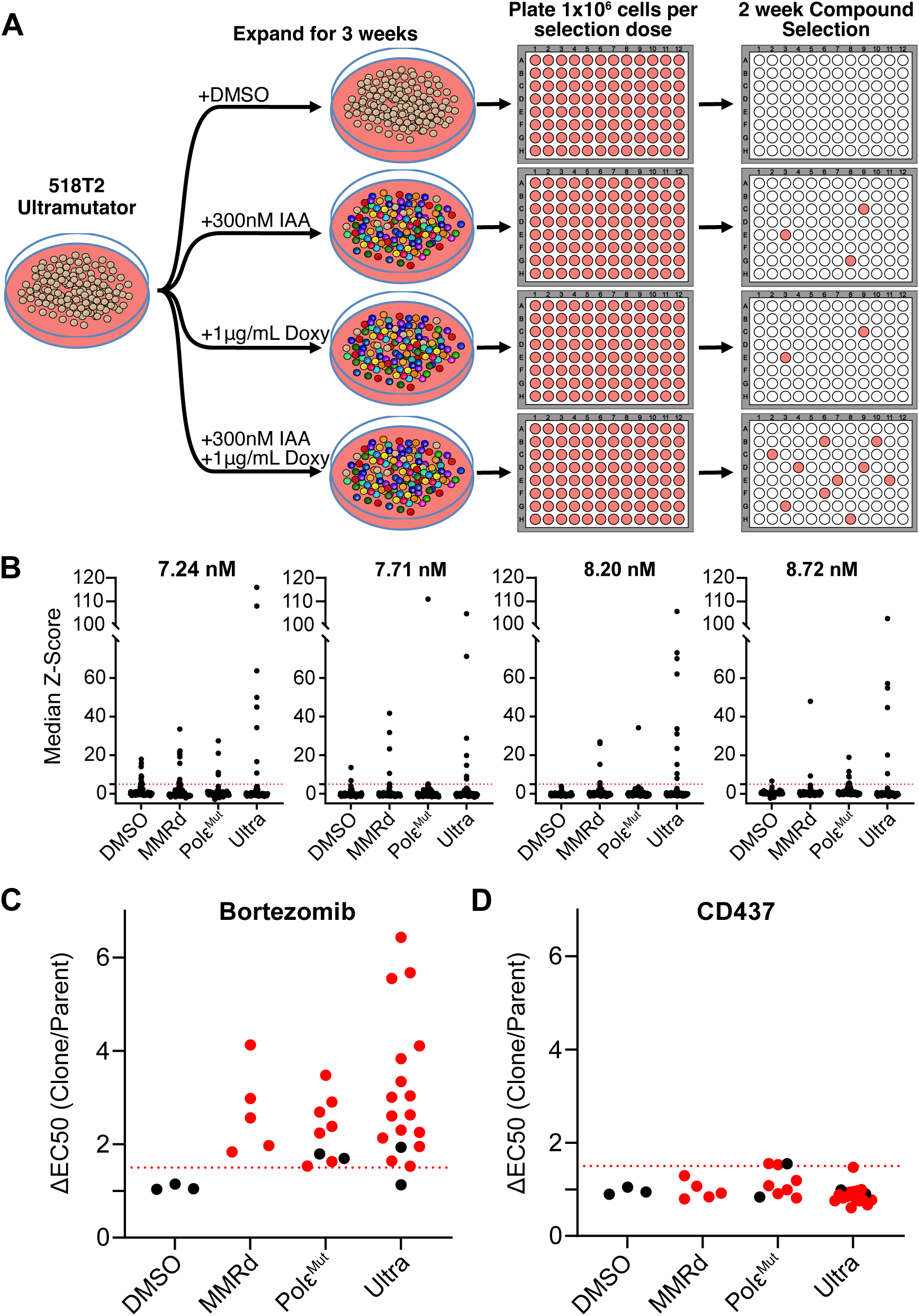
Emergence of resistant clones increases under ultramutator condition. (A) A schematic for forward genetics screening using 518T2Ultra. (B) Resazurin viability assay for 518T2Ultra cells treated with indicated concentrations of bortezomib. Red dotted line indicates Median Z-Score = 5 (C) Ratio of bortezomib EC50 of selected clones to parental 518T2Ultra, dotted red line indicates ΔEC50 = 1.5. (D) Ratio of CD437 EC50 of selected clones to parental 518T2Ultra, dotted red line indicates ΔEC50 = 1.5.

To determine whether combined MMR deficiency and *Polε^P286R^*expression (hereafter referred to as MMRd-*Polε^P286R^*) enhanced the emergence of drug-resistant clones relative to either perturbation alone, we performed proof-of-principle resistance screens using bortezomib, a proteasome inhibitor with a well-characterized target (*Psmb5*) and established resistance-conferring mutations^16, 17^. To establish selection conditions, we first determined the minimum bortezomib concentrations required to achieve complete cell killing after one week of drug exposure. We then performed resistance selections at 7.24, 7.71, 8.2, and 8.72 nM bortezomib in untreated control cells, 5-Ph-IAA treated MMR-deficient cells, doxycycline treated *Polε*^P286R^-expressing cells, and 5-Ph-IAA/doxycycline-treated MMRd-*Polε*^P286R^ cells (Supplementary Figure 3A). Across all bortezomib concentrations tested, the non-mutagenic condition (DMSO) produced the fewest resistant clones. In contrast, MMRd-*Polε^P286R^* cells consistently yielded more resistant clones than either MMR-deficient or *Polε^P286R^* expressing cells alone (Figure 3B).

Resistant clones from each condition were isolated, expanded, and subjected to dose-response analyses to confirm bortezomib resistance. To evaluate whether resistance resulted from a compound-specific mechanism rather than a generalized multidrug-resistant phenotype, we also assessed sensitivity to CD437, a DNA polymerase α inhibitor (Figure 3C-D; Supplementary Figure 3B-I). Clones isolated from the DMSO control condition exhibited bortezomib sensitivities comparable to those of parental Msh2-AID;TIR1^F74A^ murine SCLC cells (EC_50_ fold-change <1.5), indicating that these colonies did not represent bona fide resistant clones. In contrast, all clones derived from MMR-deficient or *Polε*^P286R^-expressing cells were resistant to bortezomib while remaining sensitive to CD437 (EC_50_ ≤ 1.5-fold higher than parental cells). Similarly, 17 of 18 clones isolated from the MMRd-*Polε*^P286R^ condition exhibited robust bortezomib resistance and retained sensitivity to CD437 (Figure 3C-J).

To identify the genetic basis of resistance, we performed Sanger sequencing of *Psmb5*. Twenty-eight of 32 resistant clones harbored previously described resistance-associated mutations in Psmb5 (Supplementary Figure 3L), including 7/9 clones derived from the *Polε*^P286R^ condition and 16/18 derived from the MMRd-*Polε^P286R^*condition^4, 5, 18^. Notably, we identified the Psmb5^C133W^ mutation, a known resistance allele previously reported in chemical mutagenesis screens that has not been observed in screens using MMR-deficient cells ^4^. Together, these findings demonstrate that combining Msh2 depletion with expression of proofreading-deficient Polε significantly enhances the emergence of bortezomib-resistant clones. The high frequency of recurrent *Psmb5* mutations supports the notion that the increased frequency of resistant clones in the MMRd-*Polε^P286R^* condition was driven by enhanced mutagenesis of the ultramutator system. We next applied the ultramutator platform to investigate mechanisms of resistance to lurbinectedin, a DNA-damaging agent approved for the treatment of extensive-stage SCLC as maintenance therapy following first-line treatment^19^. Lurbinectedin binds covalently to guanine residues within GC-rich regions of the DNA minor groove, generating DNA adducts that are processed by the transcription-coupled nucleotide excision repair (TC-NER) pathway. During this repair process, TC-NER-mediated excision of lurbinectedin-DNA adducts results in the formation of DNA double-strand breaks, leading to activation of the DNA damage response and ultimately apoptosis.^20–23^ To identify genetic determinants of lurbinectedin resistance, we performed forward genetic screens using five lethal concentrations of lurbinectedin in ultramutator (300 nM 5-Ph-IAA and 1 μg/mL doxycycline) and mutagenesis-off control (DMSO) Msh2-AID;TIR1^F74A^ murine SCLC cells. Following selection, resistant clones were isolated and expanded for further characterization Across selections performed at lurbinectedin concentrations ranging from 161 to 259 pM, we recovered nine resistant clones (Figure 4A), seven of which were determined to be independent based on unique DNA barcodes introduced prior to selection (Lurb-A through Lurb-G). Dose-response analyses confirmed that all seven independent clones were resistant to lurbinectedin, exhibiting greater than two-fold increase in EC50 relative to the DMSO-treated parental ultramutator line. Importantly, these clones retained sensitivity to bortezomib, indicating that resistance was specific to lurbinectedin rather than the result of a generalized drug-resistant phenotype (Figure 4B-D).

**Figure 4.**
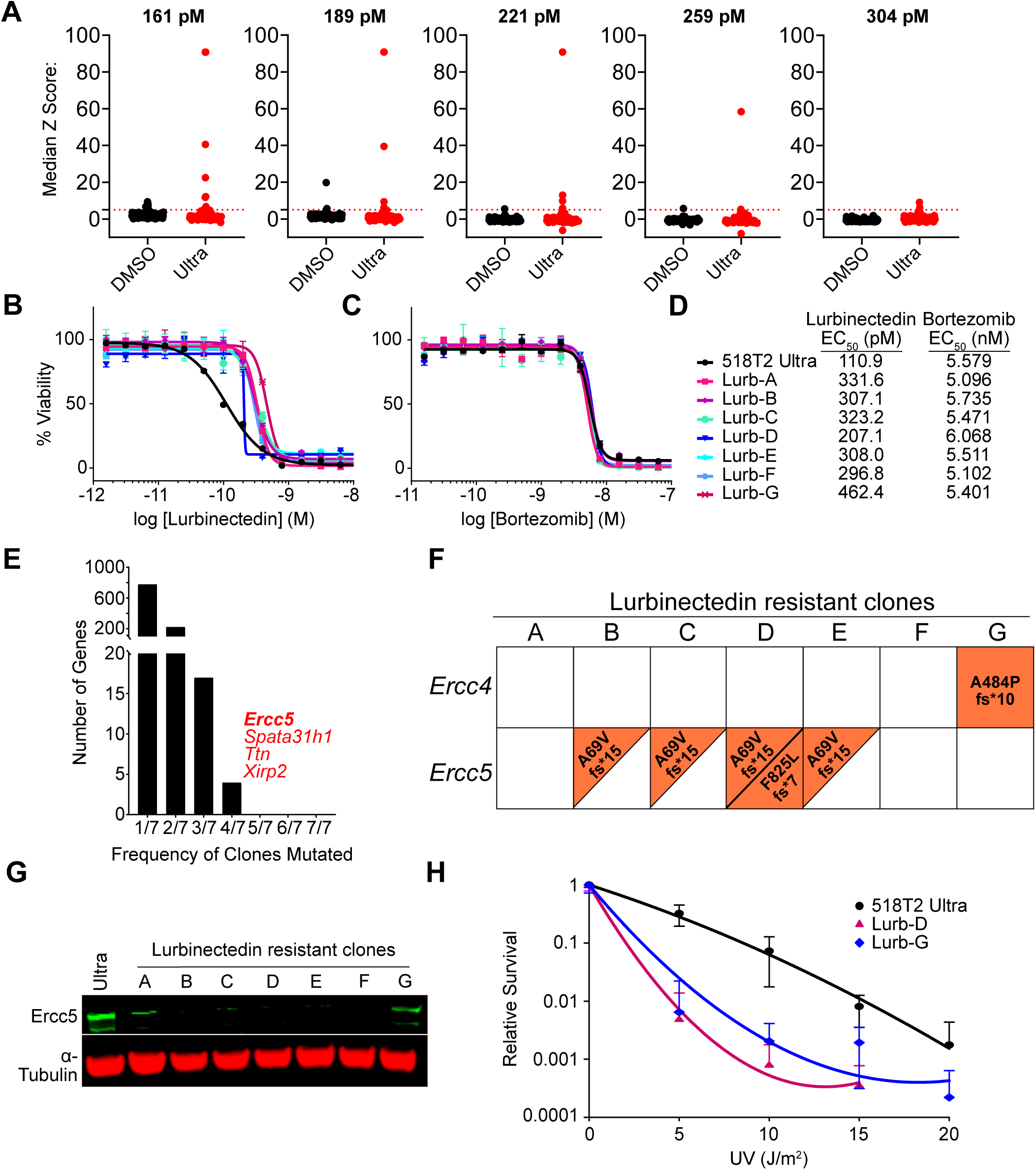
Forward genetics screening using 518T2Ultra reveals the mechanism of resistance for lurbinectedin. (A) Resazurin viability assay for 518T2Ultra cells treated with indicated concentrations of lurbinectedin. Median Z-Score for wells with oversaturated flourescence were calculated with RFU=100,000 (B) CellTiter-Glo Viability assay for lurbinectedin-resistant clones treated with lurbinectedin for 3 days. (C) CellTiter-Glo Viability assay for lurbinectedin-resistant clones treated with bortezomib for 3 days. (D) EC50 for lurbinectedin and bortezomib for each lurbinectedin-resistant clone. (E) Ercc5 is mutated in 4 out of 7 clones. (F) WES analysis results for lurbinectedin-resistant clones reveals mutations in ERCC4 and ERCC5. (G) Western blot of lurbinectedin-resistant clones for Ercc5. α-Tubulin was used as a loading control. (H) Clonogenic survivability assay measures radiation sensitivity of 518T2Ultra, Lurb-D, and Lurb-G.

Whole-exome sequencing of lurbinectedin-resistant clones (Lurb-A through Lurb-F) identified recurrent mutations in four genes (*Ercc5*, *Spata31h1*, *Ttn*, and *Xirp*) (Figure 4E). Among these candidates, *Ercc5* was particularly notable, as multiple clones harbored frameshift mutations predicted to result in loss of function. This finding is consistent with previous studies demonstrating that ERCC5 deficiency conferred resistance to lurbinectedin (Figure 4F).^22^ To determine whether these genetic alterations affected Ercc5 protein expression, we performed immunoblot analysis. Clones Lurb-B through Lurb-F exhibited complete loss of detectable Ercc5 protein, whereas Lurb-A displayed substantially reduced expression (Figure 4G). Interestingly, no coding mutations in *Ercc5* were identified in Lurb-A or Lurb-F despite diminished protein expression, suggesting that non-coding alterations may have driven the loss of Ercc5 in these clones. In addition, Lurb-B, Lurb-C, and Lurb-E each harbored a frameshift mutation in only one *Ercc5* allele, while the second allele appeared wild type by exome sequencing. Despite this, Ercc5 protein was undetectable in all three clones (Figure 4G). In contrast, Lurb-D contained two independent *Ercc5* mutations, consistent with the observed complete loss of protein expression. Together, these findings suggested that additional genetic alterations, potentially located in non-coding regulatory regions or structural variants not captured by exome sequencing, contributed to reduced or absent *Ercc5* expression observed in lurbinectedin-resistant clones, including Lurb-A, Lurb-B, Lurb-C, Lurb-E, and Lurb-F.

Reduced or absent *Ercc5* expression accounted for lurbinectedin resistance in all resistant clones except Lurb-G, which retained ERCC5 expression comparable to that of parental cells. Notably, whole-exome sequencing revealed a homozygous frameshift mutation in *Ercc4* in the Lurb-G clone (Figure 4F). *ERCC4* encodes XPF, a critical component of the nucleotide excision repair (NER) pathway. Loss of several NER factors, including XPG (ERCC5), XPF (ERCC4), XPA, and XPD (ERCC2), was previously shown to confer resistance to ecteinascidin-743, the parent compound of lurbinectedin.^22^ These observations suggested that Lurb-G also harbored an NER defect despite retention of ERCC5 expression.

Because suitable antibodies for murine Ercc4 were unavailable, we sought to functionally assess DNA repair capacity in Lurb-G cells. Defects in NER are known to confer hypersensitivity to DNA-damaging agents, including ultraviolet (UV) radiation, due to impaired repair of bulky DNA lesions.^24^ Therefore, we compared the sensitivity of parental 518T2 Msh2-AID;TIR1^F74A^ murine SCLC cells, the ERCC5-deficient clone Lurb-D, and the *Ercc4*-mutant clone Lurb-G following exposure to increasing doses of UV radiation. After irradiation, cells were allowed to recover and viability was assessed. Both Lurb-D and Lurb-G exhibited markedly increased sensitivity to UV radiation relative to parental 518T2 Msh2-AID;TIR1^F74A^ cells, consistent with impaired NER function (Figure 4G-H). Together, these findings demonstrated that the lurbinectedin-resistant clones identified in our screen harbored defects in NER, arising through loss-of-function alterations in either *Ercc5* or *Ercc4*. These results further supported the central role of NER in mediating the cytotoxic effects of lurbinectedin.

## DISCUSSION

Here, we engineered an inducible ultramutator system in cancer cell lines by combining MMR deficiency through Msh2 degradation with expression of a proofreading-deficient *Polε* mutant. Simultaneous induction of *Polε* mutant and Msh2 degradation resulted in a marked increase in the emergence of bortezomib-resistant clones relative to either MMR deficiency or mutant Polε expression alone, consistent with an elevated mutation rate following combined loss of MMR and expression of an exonuclease-deficient DNA polymerase (*Polε*). These findings mirror observations from biallelic MMR-deficient tumors, which frequently acquire secondary mutations in *POLE* or *POLD1* and subsequently develop exceptionally high mutational burdens. ^11^ Similarly, mouse models harboring proofreading-deficient *Polε* alleles develop a broad spectrum of tumor types with elevated mutational burden, further supporting the central role of defective polymerase proofreading in driving accelerated mutagenesis and tumor evolution.^25, 26^ Together, these observations suggest that combining MMR deficiency with impaired polymerase proofreading recapitulates a naturally occurring mechanism of hypermutation and provides a powerful experimental platform for enhancing mutagenesis and genetic diversity in forward genetic screens. The inducible ultramutagenesis observed in our system enabled the identification of resistance-associated mutations to clinically relevant compounds including lurbinectedin. Similar forward genetic approaches have previously been used to define mechanisms of drug resistance^2, 3, 5, 7, 10^. However, the elevated mutation rate generated by our model may allow more comprehensive sampling of mutational space, including a broader range of codons within drug targets as well as additional genes that contribute to resistance mechanisms. Supporting this idea, we identified the *Psmb5^C133W^*mutation, which had not previously been reported in MMR-deficient models but was identified in chemically induced mutagenesis screens ^4, 27^. In addition, the lurbinectedin-resistant alleles identified in this study were exclusively loss-of-function frameshift mutations, in contrast to prior reports that predominantly identified missense mutations. Moreover, the ultramutator cell line enabled the emergence of loss-of-function resistant mutations requiring biallelic inactivation, as observed in lurbinectedin clone D. This ultramutator strategy may therefore facilitate the identification of resistance mechanisms for compounds whose targets are less tractable by conventional approaches, including drugs that do not act through a single protein target. Together, these findings suggest that inducible ultramutagenesis may uncover resistance mechanisms that were inaccessible using conventional mutagenesis approaches.

Although our results do not identify a direct protein target of lurbinectedin, our results are consistent with previous studies demonstrating that an intact NER pathway is required for lurbinectedin cytotoxicity^22^. Notably, no coding mutations in *Ercc5* were detected in the Lurb-F clone despite complete loss of Ercc5 protein expression. This observation raises the possibility that resistance may arise through noncoding regulatory mutations that suppress *Ercc5* expression or through loss-of-function alterations in factors required for *Ercc5* transcriptional regulation, mRNA processing, or protein stability. Transcriptional stalling of RNA polymerase II (RNA Pol II) has been proposed as a key mediator of lurbinectedin-induced toxicity.^28^ However, we did not identify mutations in RNA Pol II subunits in any of the resistant clones. While the absence of such mutations does not exclude a role for transcriptional stalling in lurbinectedin activity, it argues against a model in which resistance commonly arises through direct alteration of RNA Pol II. Moreover, given the essential role of RNA Pol II in cellular viability, mutations that disrupt drug binding while preserving normal polymerase function may be intrinsically rare. Instead, our findings support a model in which disruption of NER represents the predominant mechanism of lurbinectedin resistance. The recurrent loss-of-function alterations identified in *Ercc5* and *Ercc4*, together with the established functional coupling between NER and transcription, suggest that impairment of NER-dependent processing of lurbinectedin-induced DNA lesions is sufficient to attenuate the cytotoxic effects of the drug.

In conclusion, we engineered an inducible ultramutator system in murine cancer cells by combining targeted degradation of Msh2 with expression of a proofreading-deficient *Polε* mutant. This approach yielded substantially more clones than MMR deficiency alone, thereby increasing the recovery of resistance-conferring mutations and facilitating the identification of drug resistance mechanisms. Our findings establish combined MMRd and proofreading deficient polymerase induction as a powerful platform for forward genetic screens aimed at elucidating drug mechanisms of action and resistance pathways.

## METHODS

### Cell Culture

Murine SCLC cell line, 518T2, was cultured at 37°C with 5% CO_2_ in DMEM (Sigma #D6429) supplemented with 5% FBS (Corning 35-010-CV) and 2mM L-glutamine (Corning 25-005-CL).

### Western Blotting

Western-blotting was performed using standard methods using pre-cast gels (Invitrogen NW04125BOX). Bio-Rad Nitrocellulose Membranes (Bio-Rad, #1620112) were used for protein transference and then blocked using Bio-Rad Blotting-Grade Blocker in TBS (Bio-Rad, #1706404) for 1 hour at RT. Primary antibodies used were anti-V5 tag (D3H8Q) (Cell Signaling Technologies #13202), anti-GAPDH (GeneTex GTX627408), anti-PolE (GeneTex GTX132100), and anti-Msh2 [D24B5] XP rabbit mAb (#2017, Cell Signaling Technology). Primary antibodies were incubated for 1 hr at RT diluted 1:1000 in Bio-Rad Blotting-Grade Blocker: TBS-Tween (0.1%). Membrane was washed with TBS-Tween (0.1%) three times for 5 minutes each wash. Secondary antibodies used were IRDye 800CW Donkey anti-Mouse (#925-32212, LI-COR), IRDye 800CW Donkey anti-Rabbit (#926-32213, LI-COR), IRDye 680RD Donkey anti-Rabbit (#926-68073, LI-COR), and IRDye 680RD Donkey anti-Mouse (#926-68072, LI-COR). Visualization was performed with Odyssey CLx Imaging System (LI-COR).

### Compounds

Bortezomib was purchased from Selleck Chemicals (#S1013). CD437 was purchased from Sigma-Aldrich (#C5865). Lurbinectedin was purchased from MedChemExpress (#HY-16293). Compounds were resuspended in DMSO (Fisher Bioreagents #BP231-100) at a concentration of 10mM in aliquots of 100 μL and kept at –20C.

### Dose-Response Curves

Cells were plated in 96-well plates, 10,000 cells per well in 200μL of media. After overnight incubation, compounds were dispensed using a D300e Digital Dispenser (TECAN). Cell viability was assessed after 72 hours using CellTiter-Glo luminescent cell viability assay (Promega, #G7571). CellTiter-Glo reagent was diluted by adding PBS-Triton-X (1%) (1:1 ratio). Luminescence was assessed using a cytation 5 plate reader (BioTek).

### Selection of Resistant Clones

Selection of bortezomib resistant clones was performed by plating cells under DMSO, 300nM 5-Ph-IAA, 1μg/mL Doxycycline, or a combination of 300nM 5-Ph-IAA and 1μg/mL Doxycycline and grown for 3wks. Cells were then plated in 96 well plates, 10,000 cells per well in 200uL of media. After overnight incubation to allow cells to adhere, bortezomib was added using a D300e Digital Dispenser (TECAN), and cells were cultured in the presence of compound for 14 days. To identify surviving clones, cells were cultured in 0.04 mg/ml resazurin (BioRad) in DMEM without phenol red, supplemented with 5% FBS, 1% penicillin/streptomycin, and 1% L-glutamine. Fluorescence was measured on a Cytation 5 plate reader (BioTek) with a 560/20 excitation filter and a 590/20 emission filter to identify wells with surviving cells. Clones identified with resazurin were verified by visual inspection, and the media of verified clones were changed to DMEM (Sigma D6429) supplemented with 5% FBS and 2mM L-glutamine. The median Z-score was determined as Median Z Score = ((Individual Well Fluorescence Intensity – Median Fluorescence Intensity per mutagenesis condition)/(Median Absolute Deviation of Individual Well Flourescence)). Surviving clones were expanded in standard growth medium.

### Identification of *Psmb5* mutations by Sanger sequencing

Genomic DNA was isolated from bortezomib and lurbinectedin resistant clones using Zymo Research Quick-DNA Miniprep (Zymo Research #D3024). The exon 2 of *Psmb5* was amplified using the gDNA from bortezomib resistant clones as template and the following primers: 5’ -AAA AAG ATG CAA AGC CAG GTT – 3’ and 5’ - GGC TCA CAG GAC ACA ACT CA – 3’. The resulting amplicon was subjected to Sanger sequencing using 5’ - GTA TGG GGA GTA TTT GTG GTC TTA CGG – 3’.

### Whole Exome Sequencing Analysis

Whole-exome sequencing of Lurbinectedin-resistant clones was performed by AZENTA Life Sciences using SureSelect Mouse All Exon V1 (mSCLC) and Illumina NovaSeq 6000. The analysis workflow was based on Genome Analysis Toolkit (GATK, v4.6.2.0) best practices^29, 30^. The quality of sequencing reads was evaluated using FastQC (v0.12.1)^31^ with aggregated reports generated by MultiQC (v1.30)^32^, and adapter trimming was performed using Trim Galore (v0.6.10)^33^. High-quality reads were mapped to the mouse reference genome GRCm39 (Ensembl release 114) using BWA-MEM2 (v2.3)^34^. GATK MarkDuplicates was used to flag PCR duplicates and GATK BaseRecalibrator was applied to recalibrate base quality scores. Variant calling was restricted to exonic regions defined by the GRCm39 Ensembl release 114 GTF annotation. Post-alignment quality metrics, including duplication rate and on-target rate, were assessed using Qualimap (v2.2.2)^35^. Somatic variant calling was performed using GATK Mutect2 in paired tumor–normal mode, with the CBC parental cell line serving as the matched normal control. Variant filtering was applied using GATK FilterMutectCalls with default parameters. Variants were annotated using Ensembl Variant Effect Predictor (VEP, v114.2)^36^ against the Mus musculus GRCm39 database. Secondary filtering was performed in R using the vcfR package (v1.15.0)^37^; somatic mutations were defined by a tumor allele frequency greater than 0.2 (VAF_mut > 0.25) and a matched normal allele frequency below 0.01 (VAF_norm < 0.01). The complete pipeline was implemented in Nextflow^38^ and executed on a SLURM high-performance computing cluster.

### Radiation treatment

For UV irradiation, cells growing exponentially on culture dishes were washed with phosphate-buffered saline (PBS) and exposed to UV-C (254 nm) at a rate of 0.4-0.5 J/m2/s to achieve the desired cumulative dose. Fresh culture medium was added to the culture dishes immediately after irradiation. The colonies were fixed and stained with 0.5% crystal violet (20% Methanol in PBS).

### Quantification and statistical analysis

Data was analyzed using Prism 11.0 by GraphPad. Fitting for dose response curves were performed by using log(inhibitor) vs. Normalized response -- Variable slope (four parameters). This was used to calculate EC_50_ of each dose response curve.

## ACKNOWLEDGEMENTS

The authors thank Florentina Vandiver for technical support. psPAX2 and pMD2.G were a gift from the McFadden Lab. XLone and pCAGE-HyPBase plasmids were a gift from Dr. Frederick Rehfeld. We thank Jeon Lee and Khoi for the support with the bioinformatic analysis. D.G.M. was supported by NIH U54CA231649 and NIH UM1CA294119. A.J.D. was supported by a Department of Energy DE-SC0025578, NIH/NINDS R01NS133439, NIH/NCI R01CA276058, and NASA 80NSSC23K1018.

## AUTHOR CONTRIBUTIONS

D.G.M. and J.M.P. conceived the study. J.M.P. and A-H.N. designed, led, and performed experiments. V.L. generated Msh2-AID;TIR1^F74A^ cells. W-M.C., A.J.D., J.M.P., and A-H.N. designed and performed UV irradiation assay, and J.K. performed PCR amplification and sanger sequencing of bortezomib-resistant clones. J.M.P. analyzed whole exome sequencing results. J.M.P., D.G.M., A.J.D., and A-H.N. wrote the manuscript.

---

**Supplementary Figure 1.**
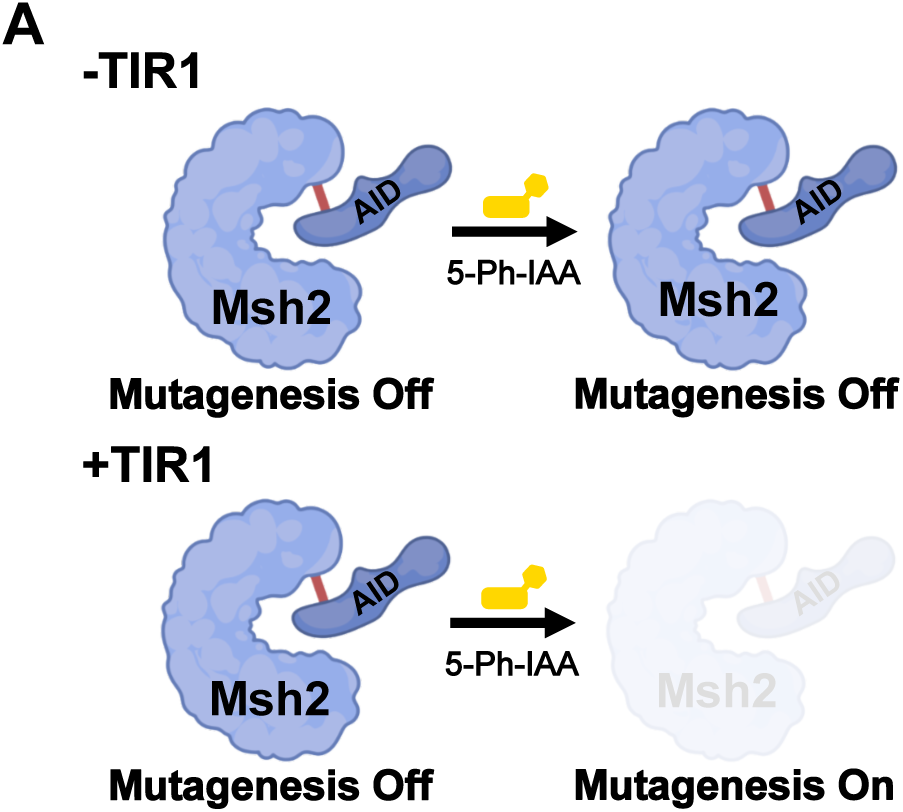
Schematic depicting the TIR1-dependent auxin inducible degradation of Msh2

**Supplemental Figure 2.**
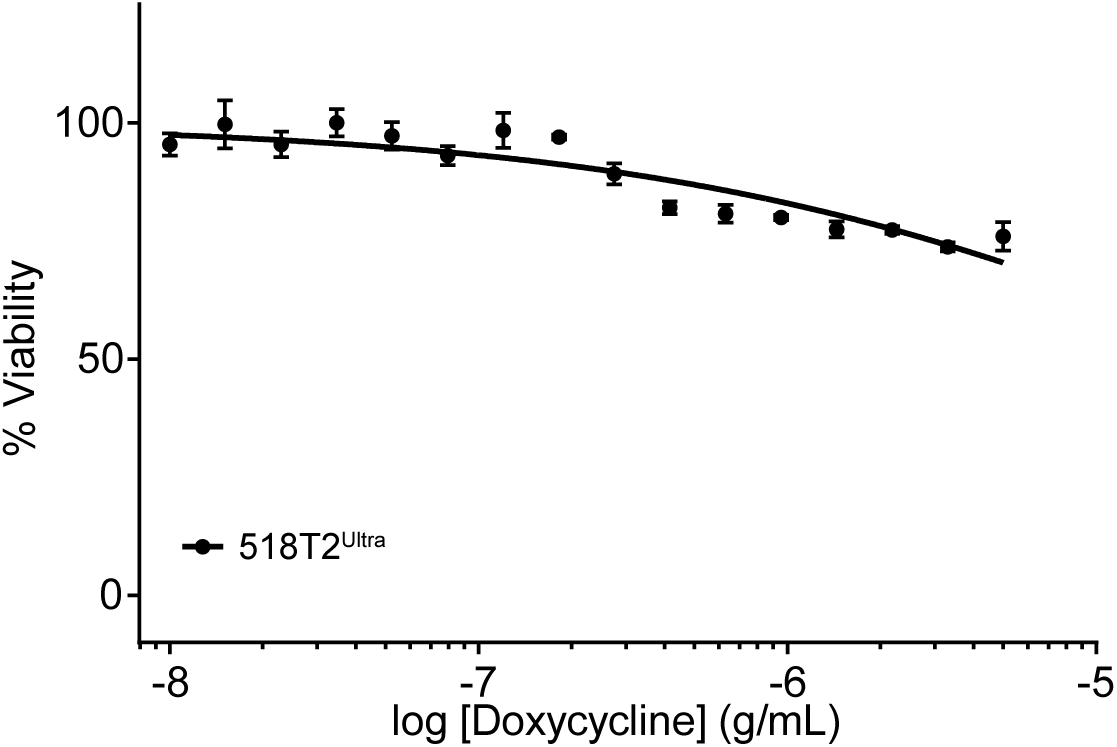
CellTiter-glo viability assay of 518T2^Ultra^ cells treated with Doxycycline for 3 days.

**Supplementary Figure 3.**
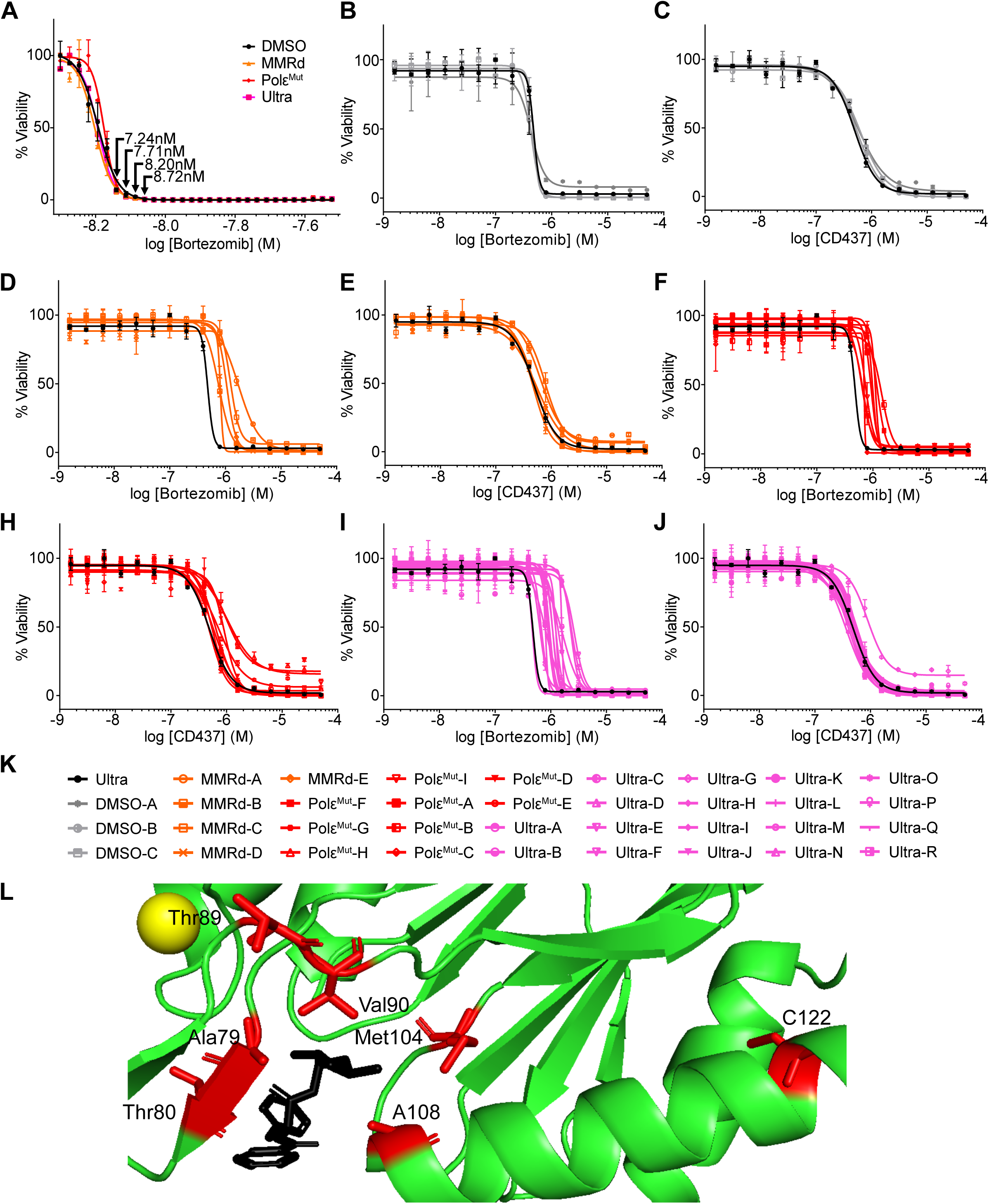
(A) Determination of bortezomib EC_100_ with indicated selection concentrations. (B-C) CellTiter-Glo viability assays with bortezomib-selected clones grown in DMSO condition and treated with Bortezomib (B) or CD437 (C) for 3 days. (D-E) CellTiter-Glo viability assays with bortezomib-selected clones grown in 5-Ph-IAA condition and treated with Bortezomib (D) or CD437 (E) for 3 days. (F-H) CellTiter-Glo viability assays with bortezomib-selected clones grown in Doxycycline condition and treated with Bortezomib (F) or CD437 (H) for 3 days. (I-J) CellTiter-Glo viability assays with bortezomib-selected clones grown in Ultramutator condition and treated with Bortezomib (I) or CD437 (J) for 3 days. (K) Legend for viability curves. (L) Crystal structure of PSMB5 bound to bortezomib (PDB: 5L5Z). Mutated residues are indicated in red.

**Supplementary Figure 4.**
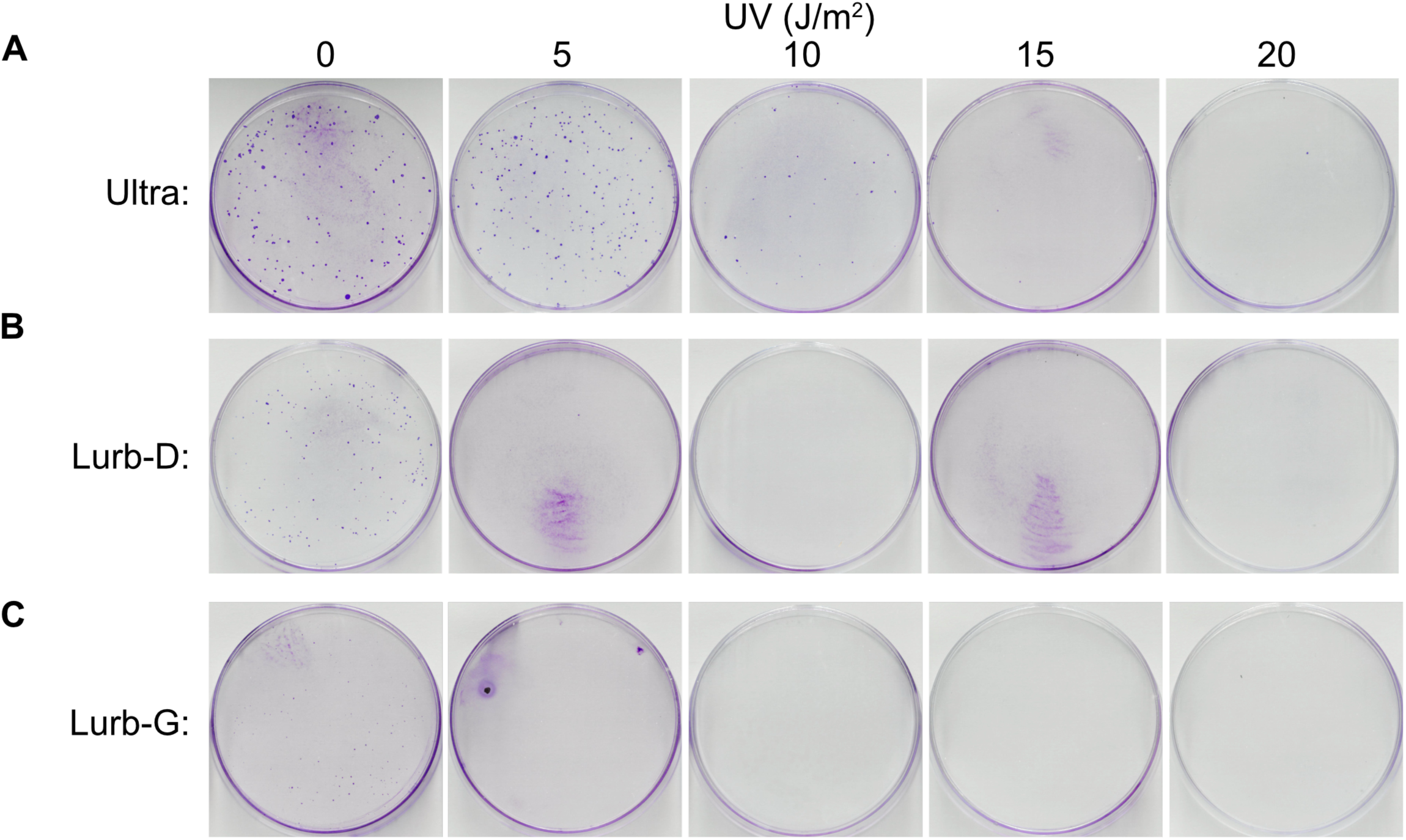
(A) Crystal violet staining of parental 518T2^Ultra^ at indicated doses of ultraviolet radiation. (B) Crystal violet staining of Lurb-D clone at indicated doses of ultraviolet radiation. (C) Crystal violet staining of Lurb-G clone at indicated doses of ultraviolet radiation.

## Notes

### Competing Interest Statement

The authors have declared no competing interest.

